# Description of Bacterial RNA Transcripts Detected in *Mycobacterium tuberculosis* – Infected Cells from Peripheral Human Granulomas using Single Cell RNA Sequencing

**DOI:** 10.1101/2024.08.20.608852

**Authors:** Philip J. Moos, Allison F. Carey, Jacklyn Joseph, Stephanie Kialo, Joe Norrie, Julie M. Moyarelce, Anthony Amof, Hans Nogua, Albebson L. Lim, Louis R. Barrows

## Abstract

*Mycobacterium tuberculosis* (Mtb) remains a global human health threat and a significant cause of human morbidity and mortality. We document here the capture of Mtb transcripts in libraries designed to amplify eukaryotic mRNA. These reads are often considered spurious or nuisance and are rarely investigated. Because of early literature suggesting the possible presence of polyadenylated transcripts in Mtb RNA, we included the H37Rv Mtb reference genome when assembling scRNA seq libraries from fine needle aspirate samples from patients presenting at the TB clinic, Port Moresby General Hospital, Papua New Guinea. We used 10X Genomics single-cell RNA sequencing transcriptomics pipeline, which initiates mRNA amplification with poly-T primers on ∼30-micron beads designed to capture, in this case, human mRNA associated with individual cells in the clinical samples. Utilizing the 10X Genomics Cell Ranger tool to align sequencing reads, we consistently detected bacterial small and large ribosomal subunit RNA sequences (rrs and rrl, respectively) and other bacterial gene transcripts in the cell culture and patient samples. We interpret Mtb reads associated with the host cell’s unique molecular identifier (UMI) and transcriptome to indicate infection of that individual host cell. The Mtb transcripts detected showed frequent sequence variation from the reference genome, with greater than 90% of the rrs or rrl reads from many clinical samples having at least 1 sequence difference compared to the H37Rv reference genome. The data presented includes only bacterial sequences from patients with TB infections that were confirmed by the hospital pathology lab using acid-fast microscopy and/or GeneXpert analysis. The repeated, non-random nature of the sequence variations detected in Mtb rrs and rrl transcripts from multiple patients, suggests that, even though this appears to be a stochastic process, there is possibly some selective pressure that limits the types and locations of sequence variation allowed. The variation does not appear to be entirely artefactual, and it is hypothesized that it could represent an additional mechanism of adaptation to enhance bacterial fitness against host defenses or chemotherapy.

## Introduction

Next-gen sequencing has facilitated studies of bacterial genomes and uncovered pathogen variants associated with clinically relevant phenotypes such as antibiotic resistance [1]. However, these studies are primarily focused on bacteria cultured from patient tissues, and thus viable but non-cultivable bacteria are not assessed. This yields an incomplete assessment of bacteria related to the infection. We document here the capture of bacterial transcripts in libraries designed to amplify eukaryotic mRNA. These reads are often considered spurious or nuisance and are rarely investigated [2]. Here a description of *Mycobacterium tuberculosis* (Mtb) ribosomal RNA sequences detected in host human cells obtained from peripheral lymph node aspirates from patients infected with Mtb.

Mtb remains a global human health threat and a significant cause of human morbidity and mortality [3]. Drug-resistant Mtb is becoming more prevalent, and so the discovery of new agents with potential anti-tuberculin activity is important [4]. We employed single-cell RNA sequencing (scRNA seq) when assessing an *in vitro* model of Mtb infection for drug discovery we developed. Because of early literature suggesting the possible presence of polyadenylated transcripts in Mtb RNA [5], we included the H37Rv Mtb reference genome (NC_000962.3 [6]) when assembling scRNA seq libraries from Mtb-THP-1 co-culture experiments. THP-1 cells are a human monocytic cell line capable of being infected by Mtb [7–11]. The detection of infected THP-1 cells hosting Mtb transcripts in our cell culture experiments promised the ability to identify individual infected host cells in clinical samples from patients infected with Mtb and to contrast their transcriptomes to other resident cell types. This ability could provide new insight into cellular responses to infection within accessible involved tissues, such as peripheral lymph nodes.

We describe the Mtb transcripts identified using this approach here. The clinical samples were fine needle aspirate samples from patients presenting at the Central Public Health Laboratory TB clinic (CPHL), Port Moresby General Hospital, Papua New Guinea (PNG), with nodal granulomas greater than 0.8 cm in diameter by external caliper measurement [12]. We used 10X Genomics single-cell RNA sequencing (scRNA-seq) transcriptomics pipeline [12,13], which initiates mRNA amplification with poly-T primers on ∼30-micron beads designed to capture, in this case, human mRNA associated with individual cells in the clinical samples. Utilizing the 10X Genomics Cell Ranger tool to align sequencing reads [13], we consistently detected bacterial small and large ribosomal subunit RNA sequences (rrs and rrl, respectively) and other bacterial gene transcripts in the cell culture and patient samples. We interpret Mtb reads associated with the host cell’s unique molecular identifier (UMI) and transcriptome to indicate infection of that individual host cell.

The Mtb transcripts detected in the THP-1 cell/H37Ra co-culture experiments exhibited significant sequence variation compared to the reference H37Ra (NC_009525.1 [6]) genome. Results obtained in this constrained system showed that approximately one-third of the detected rrs or rrl transcripts of H37Ra exhibited nucleotide variations at one or more sites. The detection of Mtb-infected cells in the co-culture experiments was relatively low but suggested that the detection of intracellular bacterial sequences from patient samples in the 10X pipeline would likely be reliable.

The frequent Mtb transcript sequence variation observed in the co-culture experiments presaged an even higher degree of transcript variation observed in the clinical samples. Greater than 90% of the rrs or rrl reads from many clinical samples had at least 1 sequence difference compared to the H37Rv reference genome. This transcript variation in the clinical samples, combined with the highly conserved nature of bacterial rrs and rrl genes, made it impossible to confirm Mtb as the infecting organism solely based on sequence homology in several clinical samples. BLAST® (blastn [14]) searches of the aligned sequences often ranked other bacteria as better matches to the detected sequences than Mtb. Therefore, to provide confidence that the bacterial transcripts detected in the clinical samples actually arose from Mtb, the data presented here includes only bacterial sequences from patients with TB infection that were confirmed by the CPHL pathology lab using acid-fast microscopy and/or GeneXpert™ analysis [12,15]. As a result, we present data from 9 individual patients who had pathology laboratory confirmed TB infection. We appreciate that this does not exclude the possibility that other bacteria could have been present in the patient’s granuloma in addition to Mtb. Still, at least it independently confirms that Mtb was present in these samples, a standard used previously in the assessment of nodal tuberculosis granulomas [16]. Bacterial RNA reads from three of the nine FNA samples did identify Mtb strains as best matches during BLASTn searches. The repeated, non-random nature of the sequence variations detected in Mtb rrs and rrl transcripts from multiple patients, suggests that, even though this appears to be a stochastic process, there is possibly some selective pressure that limits the types and locations of sequence variation allowed. The variation does not appear to be entirely artefactual, and it is hypothesized that it could represent an additional mechanism of adaptation to enhance bacterial fitness against host defenses or chemotherapy.

## Results

### Detection of Bacterial Transcripts in THP-1 cells

We developed a flow cytometry-based system to quantify drug effects on different cellular compartments observable in THP-1/Mtb *in vitro* co-cultures, using GFP H37Ra (Fig. 1), to more fully reflect the intracellular course of Mtb infection [17–23]. We confirmed intracellular Mtb using confocal microscopy (Fig. 1). We also conducted scRNA-seq analysis on parallel cultures to seek transcriptional signatures of infection that might serve as valuable identifiers of infected cells from clinical samples (Fig. 2).

**Fig. 1.**
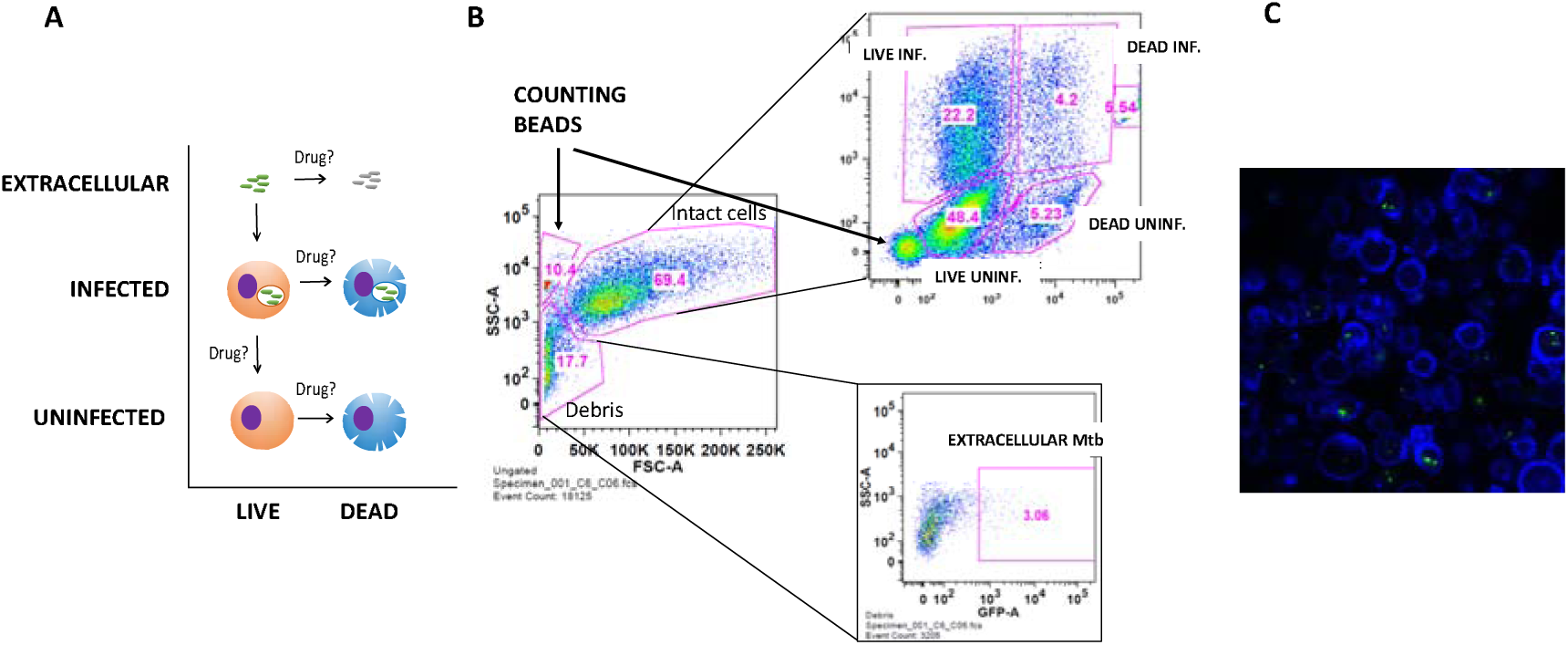
**A)** Cartoon showing THP-1 cell types quantified. THP-1 cells were differentiated with PMA for 24 h prior to infection and then co-incubated with GFP H37Ra for 5 days (MOI 2:1). **B)** Gating paradigm of intracellular and extracellular Mtb populations using forward and side scatter parameters. Internal standard counting beads quantify total cell count gain or loss. Intact cells were plotted versus V450 viability stain (abscissa) and GFP expression (ordinate). Flow cytometry conducted with a BD Canto, results analyzed using FlowJo™ software. Extracellular GFP-Mtb were in the “Debris” gate. **C)** Confocal microscopy of THP-1 cells infected with GFP H37Ra confirmed intracellular Mtb. Cells were fixed in 1% formaldehyde. Actin was stained with Abnova™ Fluorescent Dye 405-I Phalloidin (blue). The image was acquired on a Nikon A1 confocal microscope using a 60x oil lens and processed using Fiji™ software. Image is an average intensity projection of 6 z-stacks, spaced 0.5 µm apart.

**Fig. 2.**
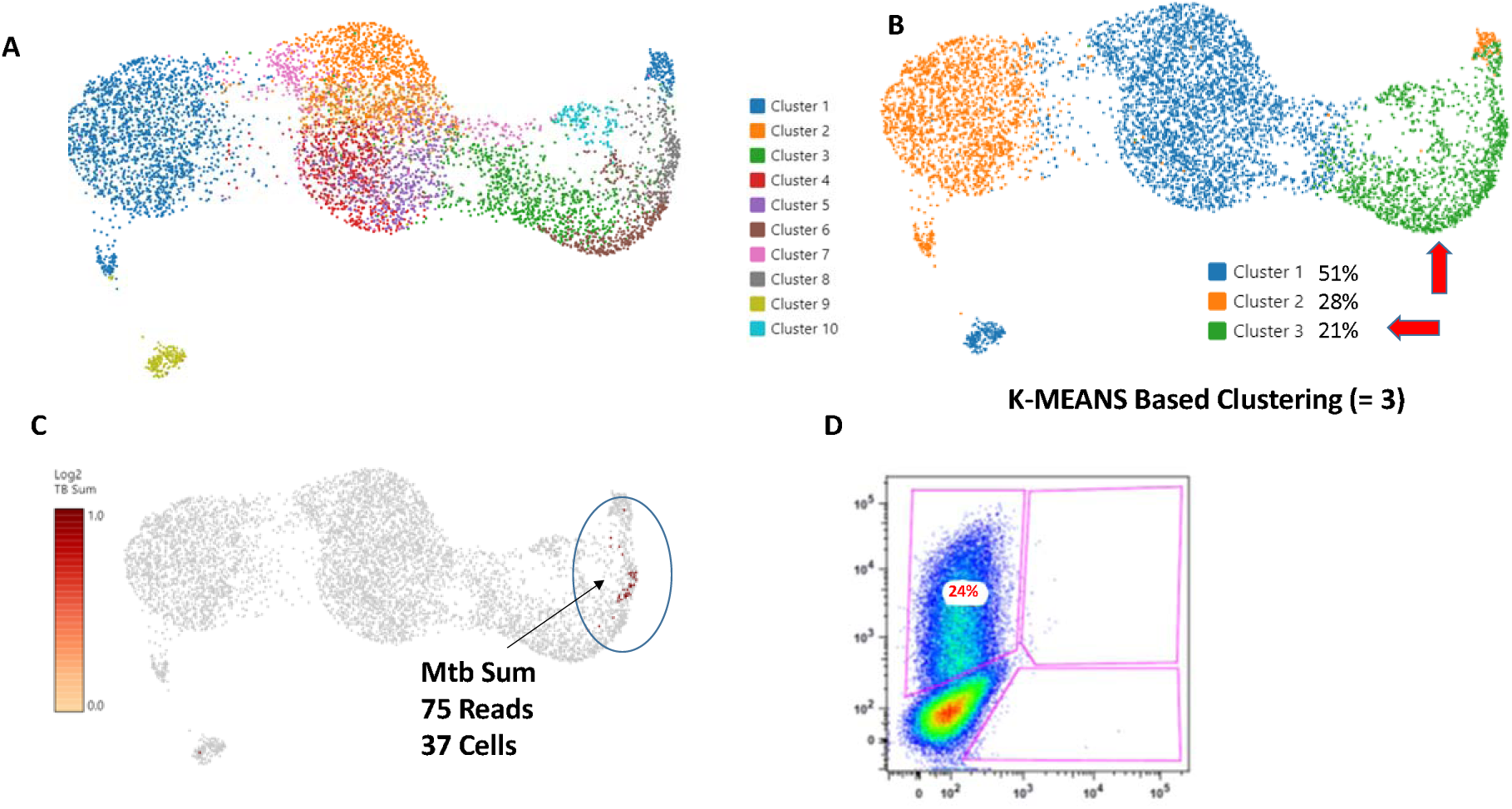
**A)** Loupe UMAP of THP-1/Mtb co-culture (14800X6) 5 days after inoculation of 0.5X10^6^ cells with GFP H37Ra. THP-1 cells were differentiated with PMA for 24 h prior to infection and then coincubated with GFP H37Ra for 5 days (MOI 2:1). **B)** Same data at K means clustering, K=3, yielded cluster 3, approximately equal to percent GFP expressing cells detected in replicate culture by Flow Cytometry. **C)** Feature plot showing any cell containing an Mtb transcript. All Mtb containing cells were found in cluster #3. **D)** Flow cytometry analysis of duplicate culture showing GFP fluorescence (percent infection) on vertical axis.

The co-culture system routinely yielded around 20% infection of THP-1 cells, defined by GFP expression, 5 days after Mtb co-culture. Replicate cultures analyzed by scRNA-seq (Fig. 2, S-1) were assessed for GFP expression using flow cytometry (Fig. 2D) and showed from 15% to 24% infection for the replicate experiments, respectively. Each replicate experiment was processed with a technical repeat, e.g., 14800X3 and - X4 and 14800X5 and -X6. Unsupervised UMAP clustering of the THP-1/Mtb co-cultures at default resolution did not yield clusters matching the percentages of GFP expressing cells determined by flow cytometry (Fig. 2A). However, K-Means based clustering at K=3 did, yielded cluster sizes almost exactly matching the percent GFP positive by flow cytometry for the given replicates (Fig. 2B, Fig. S-1). We included the Mtb H37Rv reference genome in the Cell Ranger genome alignment and queried if any Mtb sequences were associated with THP-1 UMIs (xf.25 reads) and plotted them on the feature UMAP (Fig. 2C, S-1). Thus, single-cell RNA sequencing of duplicate cultures confirmed H37Ra rrs or rrl transcripts in GFP-expressing THP-1 cells. Flow cytometry parameters were set to count a minimum of 30,000 events. An average of about 30 infected (Mtb+) cells was detected in each experiment, meaning that rrs or rrl sequences were only detected in about 3% of the GFP Mtb-infected cells. While the percentage of detected host THP-1 cells containing Mtb transcript sequences was low, all of these cells clustered in the presumed “infected clusters” determined by the percent GFP-positive cells in the parallel duplicate experiments.

Comparison of the detected Mtb sequences in repeat experiments to the reference rrs and rrl sequences (synonyms Rvnr01 and Rvnr02, respectively) in the H37Rv and H37Ra genomes (Rv and Ra are identical through the rrs and rrl genes) showed that approximately 58% (17/29) of the reads from the co-culture experiments contained at least 1 sequence variation, almost 17% of the reads (5/29) contained multiple sequence variations (Fig. 3A). These sequence variations were initially attributed to transcription errors in the 10X Genomics amplification process, but assessment of clinical samples, below, suggests that this is not a random occurrence and that additional factors may be contributing to the high read-sequence variability. The number of reads per gene across the Mtb genome was summed, and transcripts for rrs and rrl far outnumbered the other genes detected, possibly because of high transcription rates of ribosomal RNA during infection. Most other gene transcripts were detected once, while a few were detected twice, Rv0636, Rv1095, Rv1461, Rv1899c, Rv3616c, and Rv3803c (Fig. 3B). Interestingly, Rv3616c is a crucial virulence gene, and Rv3803c is a major antigen, and thus may also represent highly transcribed genes [24, 25].

**Fig. 3.**
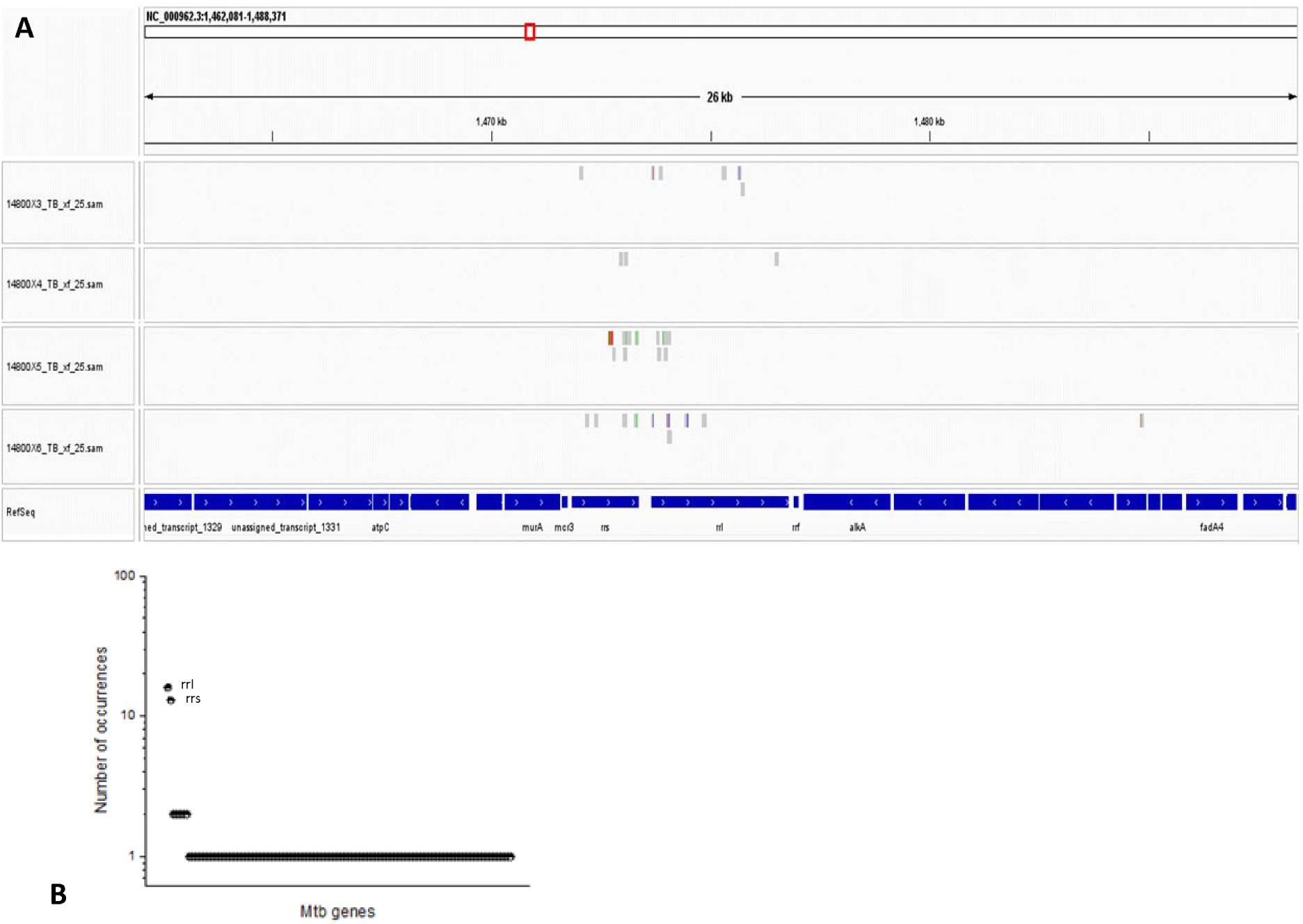
Integrated Genome Viewer alignment of reads detected in cell culture experiments with the H37Rv Mtb genome (NC_000962), centered on rrs and rrl genes. H37Rv and H37Ra (NC_009525) genomes are identical through the rrs and rrl genes. Two independent experiments, 14800X3&4 and 148005&6, were conducted. Each experiment was completed with full technical repeats designated 3 & 4 and 5 & 6 (respectively). A) Shows all Mtb reads detected throughout the rrs and rrl genes for the repeat co-culture experiments. Sequence deviations from the reference genome are indicated by color bars; changes to G (brown), C (blue), A (green), and T (Red). B) Reads of given Mtb genes detected co-culture experiments. Reads for rrl and rrs were detected most frequently. Reads for other genes with 2 copies detected were Rv0636, Rv1095, Rv1461, Rv1899c, Rv3616c and Rv3803c. The remaining transcripts detected only once totaled 130 genes.

### Coverage of the Mtb genome detected using WGS was sparse

In hoping to access a large cohort of tuberculosis patients in PNG, we sought to determine which strains of TB characterized lymph node tuberculosis (LNTB). In a trial whole genome sequencing (WGS) experiment, 5 of the original 2019-FNA samples were analyzed as a proof-of-concept for this approach.

The cell yield of methanol-preserved FNA biopsies ranged from 6.67×10^6^/mL to 3.36×10^7^/mL in ∼1.25 mL each (n = 8). After storage and transportation on ice, 0.5 mL of the cells were rehydrated following the 10X Genomics protocol [13], and the DNA was isolated for WGS. Sufficient DNA yields were obtained from 5 of 8 samples submitted for WGS analysis (75 ng DNA per sample) using Nextera Flex Technology and Illumina S4 flow cell sequencing. Samples were analyzed using standard short-read aligners, variant calling algorithms, and annotation methods [26–29]. WGS data sets for all 5 samples and analyzed for Mtb sequences.

While Mtb DNA sequences were identified in each of the analyzed samples, they were extremely rare and represented only a small fraction of the Mtb genome. Blastx results confirmed: 30S ribosomal protein S1; 50S ribosomal protein L4; DNA-directed RNA polymerase; RNA polymerase sigma factor RpoD; recombination factor protein RarA; arabinosyltransferase C and PPE family protein genes. We concluded that because the Mtb DNA was such a minor component, compared to the vast abundance of human DNA, enrichment of bacterial DNA will be necessary for comprehensive WGS of the infecting Mtb strain(s).

### Detection of Mtb transcripts in patient FNA using scRNA-seq

Bacterial transcripts were detected in 21 of the 24 patient samples [12]. The combined data set UMAP is shown in Fig. 4A. It is important to reiterate that only the results from the 9 patient FNAs that were confirmed as positive for Mtb infection by acid-fast microscopy or GeneXpert® are included here (Fig. 4B). The majority of the detected Mtb sequences mapped to rrs or rrl, as was observed in the cell culture experiments. All host cells containing any Mtb transcripts were retained in this analysis, exempting them from the more rigorous transcriptome quality control defaults applied to the rest of the cells in the respective samples.

**Fig. 4.**
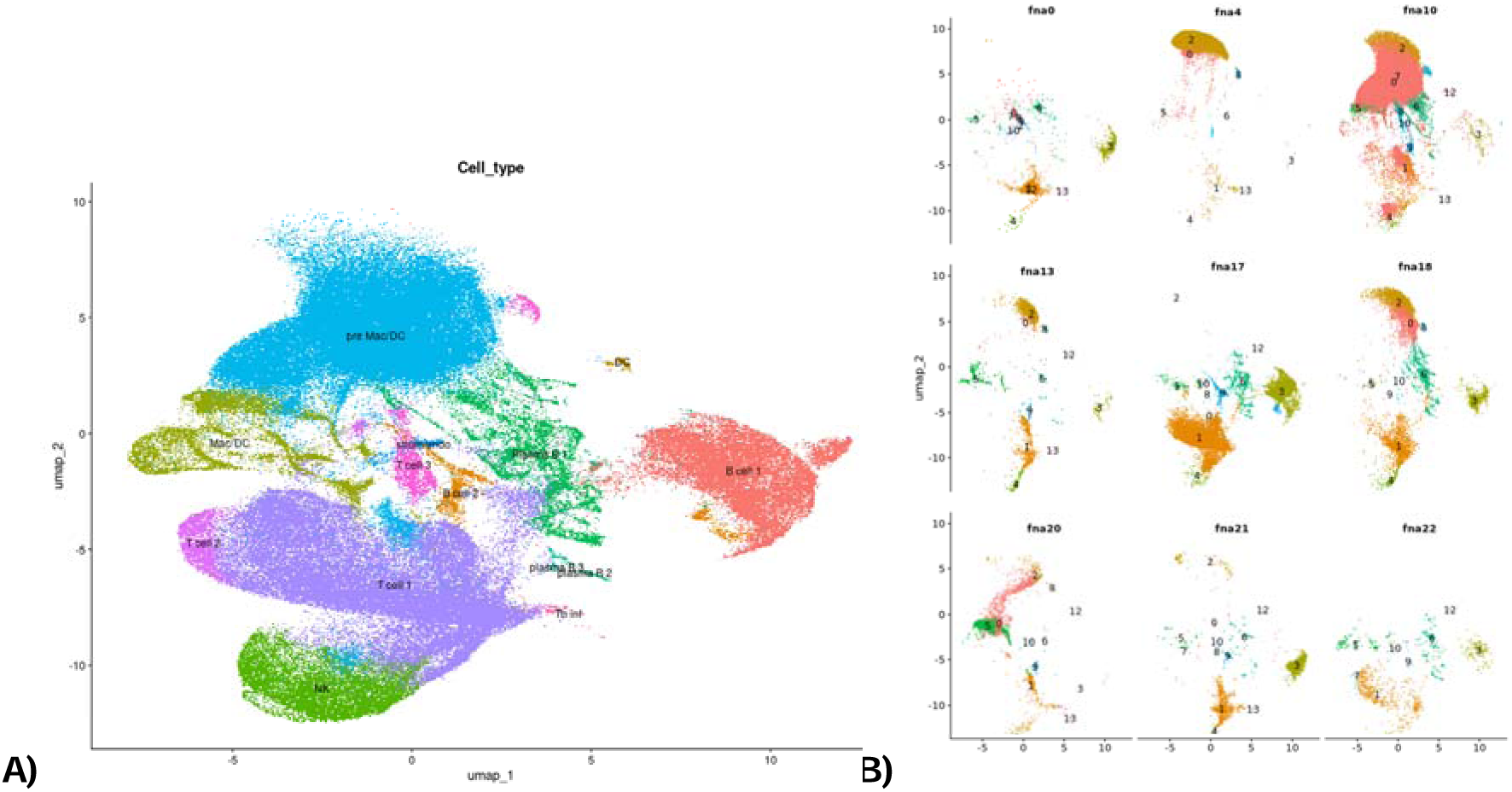
**A)** Annotated cluster identification of combined FNAs UMAP. Cluster #2 (unidentifiable cells) removed [12]. **B)** UMAPs of the 9 individual patient samples for which pathology lab confirmed the presence of Mtb in FNA samples.

### Detection of High sequence variation in Mtb Transcripts from clinical samples

Bacterial transcripts were detected in every clinical sample with confirmed TB. As in the co-culture experiments, above, bacterial rrs and/or rrl transcripts were most frequently detected. Over 90% of the transcripts in the rrs or rrl genes in most of the clinical samples contained at least one sequence difference, most of them showing multiple differences, when compared to the reference H37Rv genome. Similar sequence variation was also observed in the transcripts of other genes that were detected in the clinical samples. As discussed below, rrl and rrs are highly conserved across clinical isolates from different lineages [30], so the sequence differences seen here are not due to differences between lineages.

Figure 5 shows contrasting examples of IGV alignments of Mtb reads from 2 patient samples with relatively high levels of bacteria transcripts compared to the other FNAs analyzed. FNA IGV stacks of coverage from all reads from the 9 FNAs are presented in Fig. 5D. The frequency of detection of bacterial reads was not consistent across all patient samples. The detection of Mtb reads was low in samples FNA 17, FNA 18 and FNA 22. Whereas in FNA 0, FNA 4, FNA 10, FNA 13, FNA 20 and FNA 21 the number of detected Mtb reads was relatively high, and higher than observed in the co-culture experiments. The degree of sequence variation also differed significantly amongst the patient samples. For instance, FNA 0 displayed higher frequencies of sequence variations from the reference H37Rv genome than FNA 20, which showed very few relative to the other samples analyzed. FNA 0, FNA 4, FNA 10, FNA 13 and FNA 21 contained higher numbers of reads exhibiting repeated nucleotide variations, compared to FNA 20 and FNA 22, with FNA 10 showing a more intermediate frequency of variation.

**Fig. 5.**
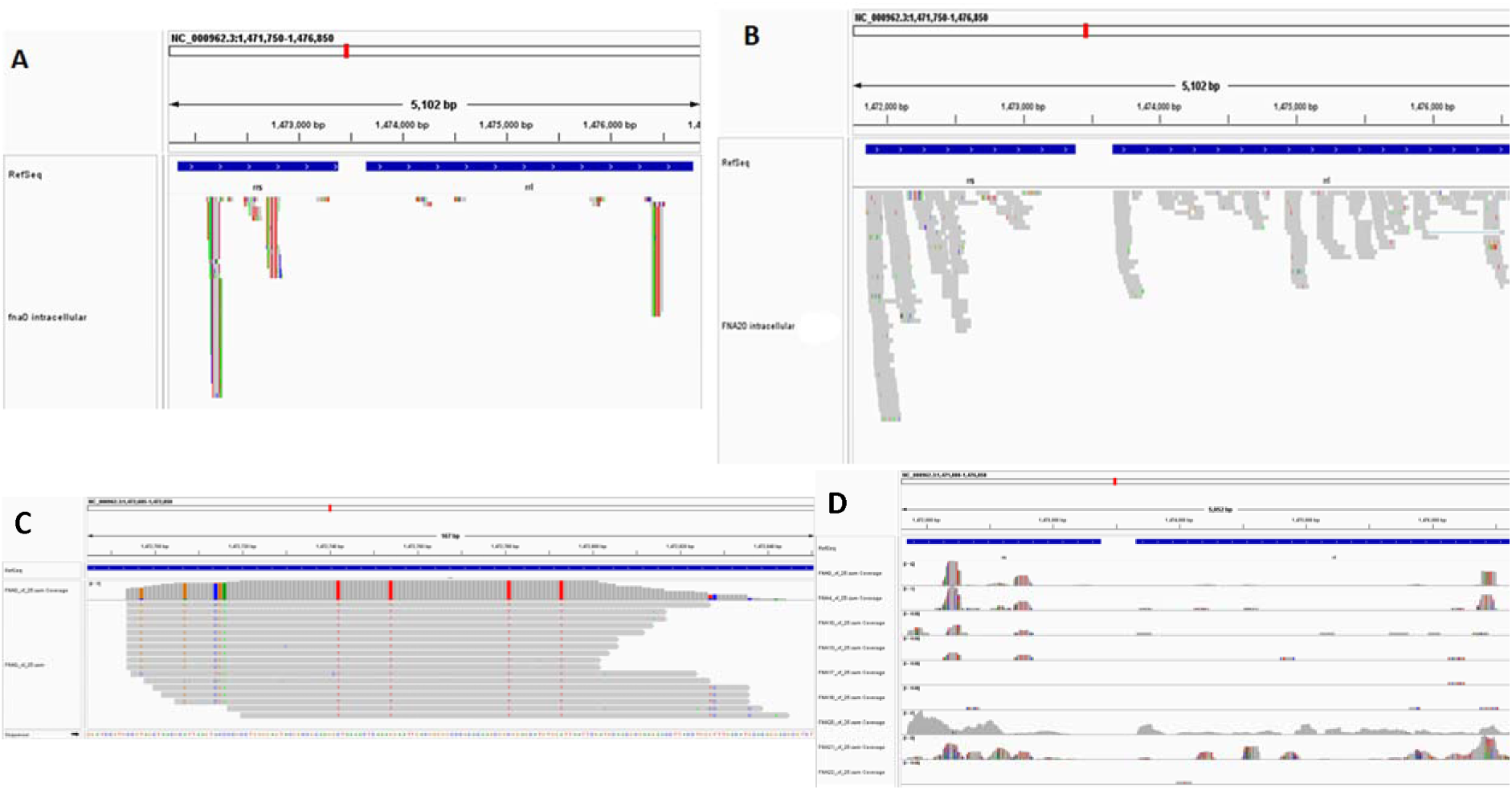
Repeated sequence deviations from the reference genome in Mtb reads. **A, B)** IGV alignment of intracellular Mtb reads from FNA 0 and FNA 20, respectively. Sequence variations versus the reference H37Rv genome are show in colors: G (brown), A (green), C (blue) and T (red). FNA 0 had a high incidence of repeated sequence variations, FNA 20 had a relative low incidence of sequence variations. **C)** A close up view of a stack of reads from FNA 0 shows sporadic sequence variations observable in independent reads (attributed to random errors in the scRNA seq pipeline). Multiple reads of identical length, starting position and directionality, usually in pairs or sets of three, are interpreted as arising from a single RNA source which we hypothesize was then amplified during library generation. Repeated/retained sequence variations are also detectable, indicated by their repetition in multiple stacked reads in this view. **D)** All intracellular Mtb reads from the 9 Mtb-confirmed FNA samples presented here in a coverage plot. Even though the samples were gathered from 9 different individuals, 1 in 2019, 1 in 2022 and 7 in 2023, the sites and types of sequence variation are repeated amongst individual infections, suggesting that the process is not random and is possibly constrained to “hard to map” gene regions or by the functionality of the changed sequence. Patterns of variation overlap and repeat in patient samples.

Closer inspection of the transcript variations revealed several details about the detected reads. First, many single nucleotide variations were repeated in multiple reads, appearing to be a retained feature, rather than a random event. We hypothesize that identical read length and directionality of the xf.25 reads indicated that the reads were replicated amplification products of the 10X Genomics pipeline. Reads of identical length almost always exhibit the same commonly repeated sequence variations. We attribute the random single nucleotide variations in individual FASTQ xf.25 reads sequences of identical length to stochastic errors introduced by the 10X Genomics sequencing process (Fig. 5C). If this interpretation is correct, then the repeated variations detected over multiple transcripts (reads of different lengths and of opposite reading orientations) would seem to suggest that the variation arises from a repeated error-prone process during library generation. Alternatively, it could also indicate some selection pressure during transcription, resulting in only certain variations. We do not believe the RNA sequence variations reflect bacterial DNA mutation because of the consistency in the published Mtb lineage WGS sequences for rrs and rrl [30]. Those WGS sequences are often derived from sub-cloned bacteria cultured from sputum samples, and thus would be expected to show diversity similar to our detected RNA reads if the diversity arose from the genome level.

Mtb transcript sequences that are associated with different UMIs clearly represent different source bacteria, although those from the same FNA might be attributed to a single inoculum. Nucleotide variation is conserved in overlapping RNA reads from bacteria associated with different UMIs (Fig. 6). The repetition of similar Mtb transcript variations associated with different UMIs shows that some transcript nucleotide variations occur repeatedly, even in different host cells. Comparison of overlapping reads from two different patient samples, also shows repeated transcript sequence variations (Fig. 6B). Showing that the repeated variations do not arise from a single inoculum of a given patient.

**Fig. 6.**
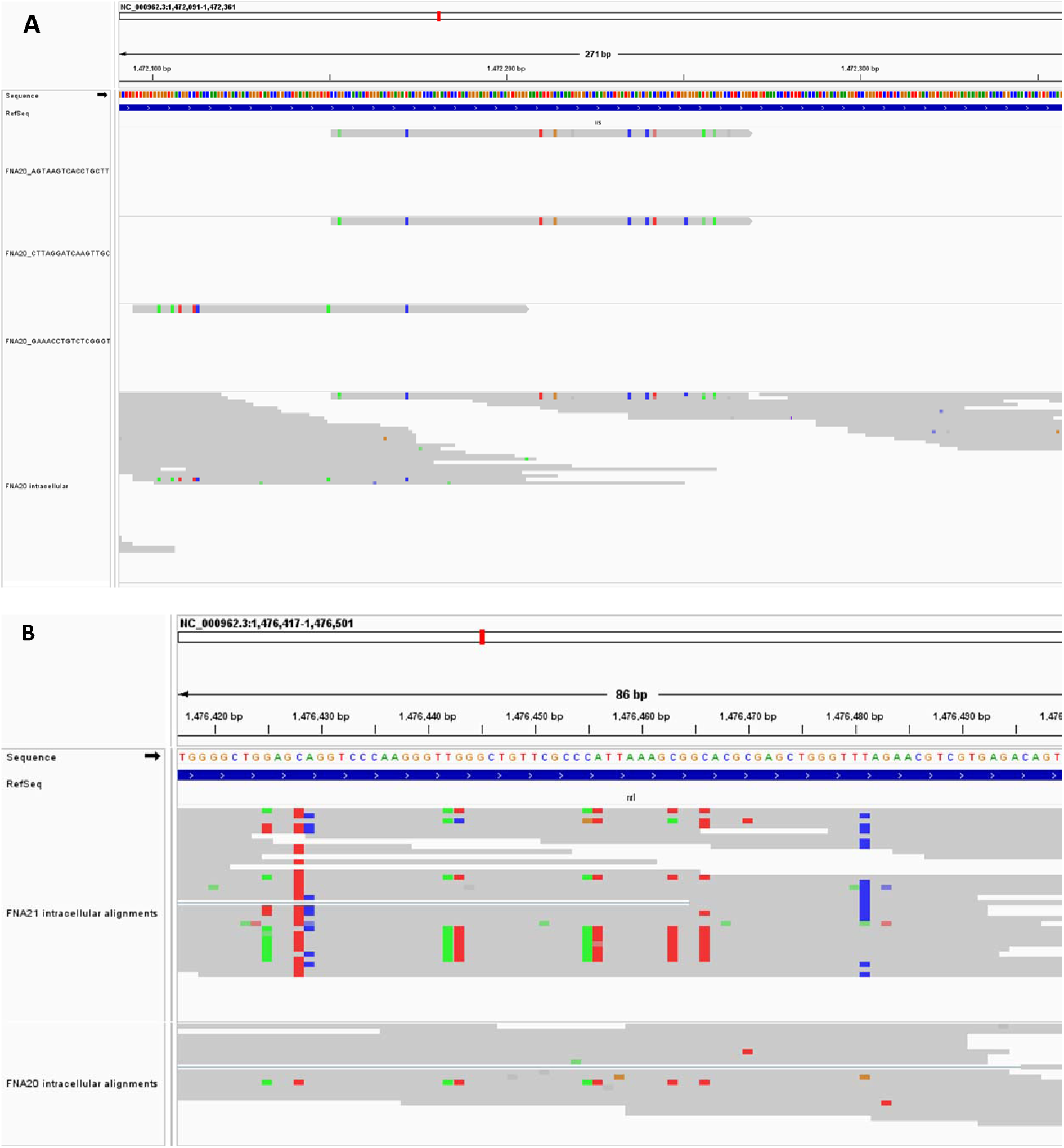
**A)** IGV view of intracellular reads from FNA 20 associated with different host cell UMIs. Overlapping reads from different host cells, and thus different infecting bacteria, retain/repeat nucleotide polymorphisms (grey bars are lower quality sequence calls). **B)** Overlapping Mtb transcript reads from two different patient samples also show similar patterns of sequence deviations from reference genome.

We attempted to compare reads from highly transcribed genes of the host human cells to see if a similar pattern of repeated transcript nucleotide variation was discernable. None of our samples had sufficient coverage of human ribosomal RNA genes for direct comparison. We did find the coverage of a highly transcribed mitochondrial (MT) gene transcript to provide sufficient overlap for comparison (Fig. 7). However, this comparison was not optimal because MT transcripts are not expected to have the extensive secondary structure and modified nucleotides found in rRNA that might represent “hard to map” or error-prone transcription-amplification sequences.

**Fig. 7.**
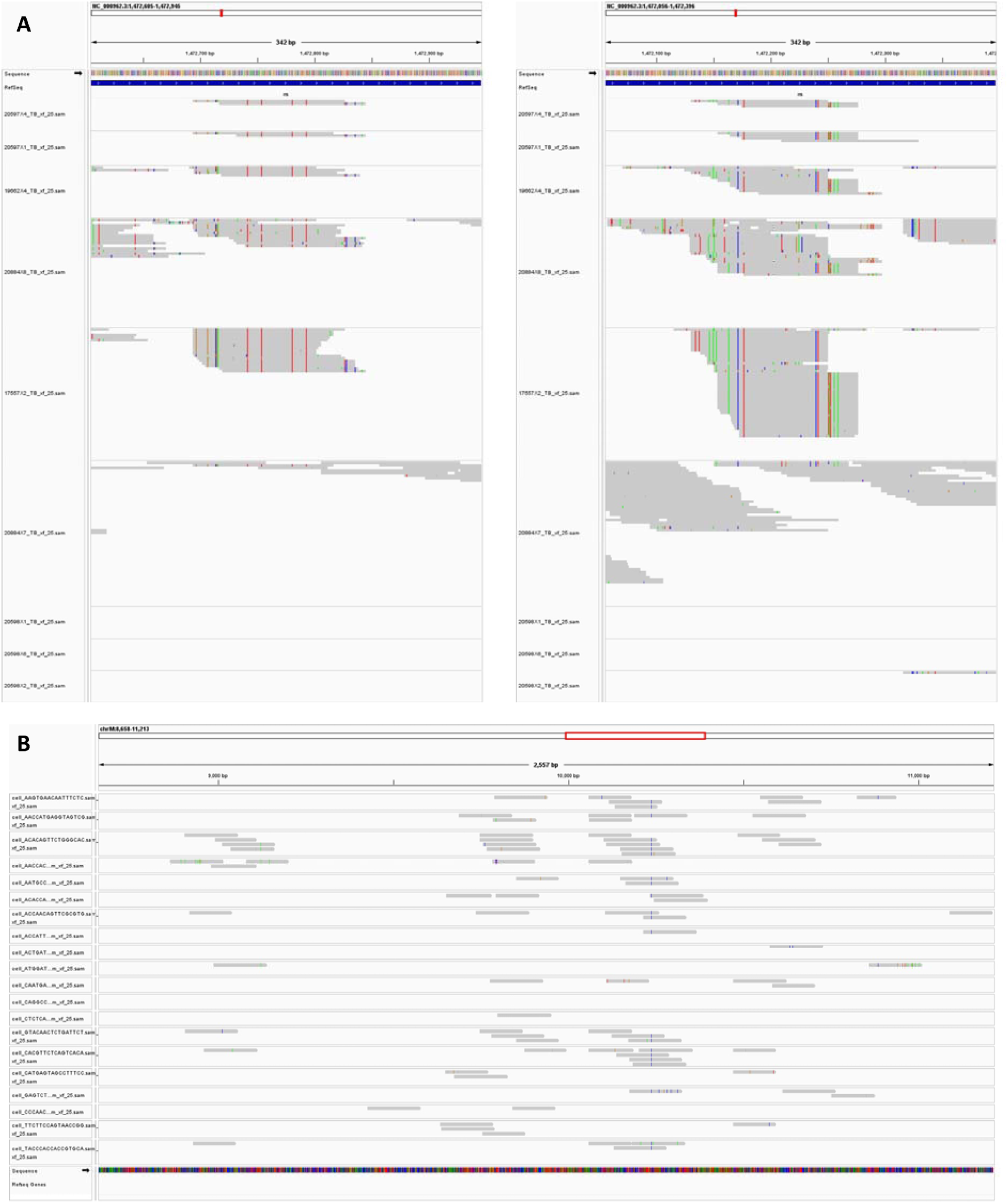
Deviations from reference sequence of Mtb reads compared to deviations found in host cell mitochondrial gene sequences. **A)** Two gene regions with overlapping Mtb reads from at least 6 of the 9 examined FNAs providing examples of repeated sequence deviations from the reference genome. FNA sequence from top to bottom is FNA 0, 4, 10, 13, 17, 18, 20, 21, 22. B) Overlapping mitochondrial gene reads from FNA 20 (top 10 cells) and FNA 21 (bottom 10 cells) single cells. All of these individual cells were also hosting Mtb sequences. Numerous random errors are seen, as well as two sequence variations repeated in every cell analyzed.

When we examined MT transcripts from individual host cells from FNA 22, we found two classes of sequence errors. One class was the expected random sequence errors expected from the 10X Genomics pipeline, errors which did not repeat in the overlapping MT reads. The second class of sequence variation was an infrequent but omnipresent change, detected in every overlapping transcript. These were observed at positions 9,123 and 10,238 in the MT gene, representing G to A and T to C transition mutations, respectively (Fig. 7B). Because of the omnipresence of these changes, we hypothesize that these could represent genomic mutations characteristic of the MT gene in our individual patient. Alternatively, these could represent hard to map nucleotides in regions of secondary structure in the MT gene that yield a consistent misreading. This second class of sequence change in the mitochondrial transcripts is similar to the repeated variations we observe in the MT gene transcripts, but it differs in its uniform consistency. In contrast, the repeated variations detected in the Mtb reads were not present in all reads without exception. We cannot rule out the possibility that human host cell rRNA reads would exhibit deviations from the parental genome sequence similar to that observed with the bacterial rRNA genes studied here, but, again, we do not have adequate coverage of the host cell rRNA to make that direct comparison.

### Some reads confirm as Mtb by BLAST analysis

When general BLAST (blastn) searches were conducted with the FASTQ rrs and rrl sequences from different patients, Mtb was the top match in only 3 of the 9 samples presented here (FNA 10, FNA 20 and FNA 22), even though all 9 FNAs were confirmed as Mtb positive by acid fast microscopy or GeneXpert® analysis. The BLAST searches were performed using three random independent batches of 5 .sam file FASTQ reads from an individual FNA. Various other bacteria were frequently identified as best matches, often with high sequence identity. However, even these “best” matches were found to vary sometimes, depending on which sets of 5 FASTQ sequences searched at a time. When this same BLAST analysis was conducted using the reads from the cell co-culture experiments, only Mtb of one strain or another was always the best match.

In a further attempt to assign our FASTQ reads to a known Mtb lineage, and because H37Rv is not the Mtb strain predominant in PNG [26, 31], we compared rrs and rrl sequences from the published WGS genomes in the reference set of clinical samples recommended by Borrell et al. [30]. Overall, there is very high identity between the published WGS data sets for these two genes across all lineages, and it was not possible to assign lineage simply using the respective published rrs and rrl sequences. Thus, we found it was impossible to use the bacterial transcripts of rrs or rrl detected by the 10X Genomics scRNA-seq pipeline to assign lineage identity to our infecting Mtb.

This was disappointing, because we found the sensitivity of this pipeline for the detection of bacterial reads rivals PCR. Several PCR attempts were made to detect transcripts of Rv1467c, RRDR, and other Mtb genes in our source re-hydrated cells, with no consistent success. This could allow for assignment of Mtb lineage in future experiments if improved coverage of the whole Mtb transcriptome is achieved.

### Detection of non-ribosomal Mtb transcripts in clinical samples

Rrs and rrl were the most frequently detected Mtb RNAs in the clinical samples (Fig. 8), with rrl and rrs being detected in 342 and 304 cells, respectively. However, these transcripts were not detected in all host cells. Many of the host cells did not exhibit detected rrs or rrl. Other gene transcripts detected included TB-Rv2186c that was detected in 6 cells. The transcript for TB-Rv2553c was detected in 5 cells; TB-Rv3343c, TB-Rv2490c, TB-Rv2319c andTB-Rv0895 were each detected in 4 cells.

**Fig. 8.**
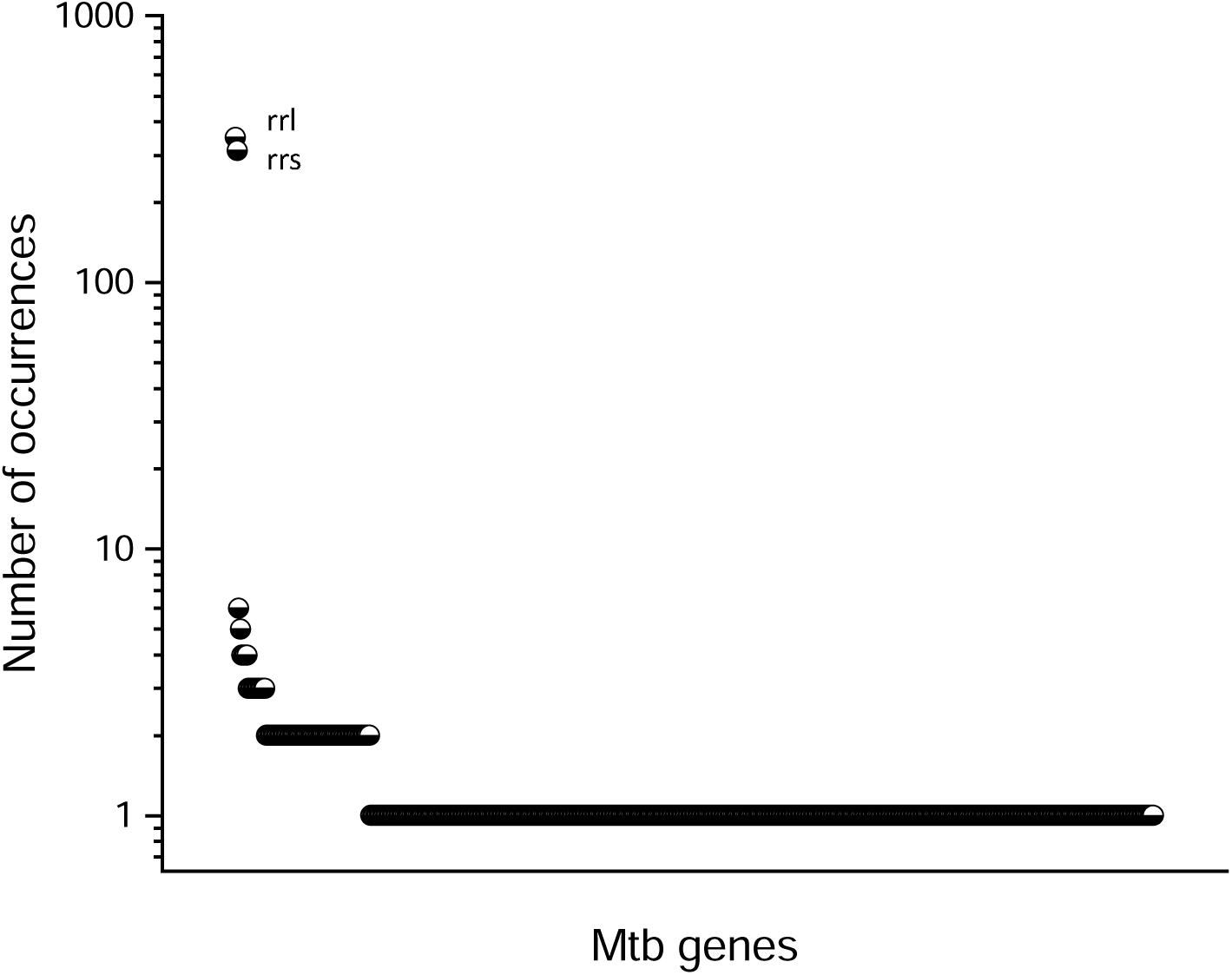
Distribution of detected Mtb RNAs throughout host cells of combined FNA samples. Plot showing frequency of Mtb transcript detection in combined Mtb positive FNAs. Transcripts for rrl and rrs were detected in 342 and 304 cells, respectively. Other gene transcripts detected included TBRv2186c that was detected in 6 cells. A total of 564 different Mtb gene transcripts were detected in the combined data set.

Eleven other Mtb transcripts were each detected in 3 individual cells, 84 different Mtb gene transcripts were detected in 2 cells, and 480 other Mtb gene transcripts were detected in only 1 cell. A total of 564 different Mtb gene transcripts were detected in the combined data set.

## Discussion

We present here the description of Mtb RNA sequences detected by single cell transcriptomic analysis of fine needle nodal granuloma aspirate samples from TB patients. Even though the 10X Genomics single cell RNA sequencing pipeline [13] is designed to capture eukaryotic mRNA, bacterial RNA sequences were detected in 21 of the 24 clinical samples. Cell culture experiments using THP-1 cells and GFP-expressing H37Ra Mtb showed that detection of Mtb rrs and rrl transcripts associated with host cell UMIs was reliable but low efficiency, estimated at approximately 3%. In scRNA-seq analysis of 12 bacteria-free THP-1 cell cultures (each conducted with a technical repeat), only 1 read was ever detected that identified as an Mtb sequence (not rrs or rrl) showing the remarkable selectivity of this analytical tool. We attribute that sole outlier to a sequencing errors inherent to 10X Genomics pipeline, and it did not occur in its technical repeat. The ability to detect individually infected cells, cluster them and compare their transcriptomes to uninfected cells in this co-culture system suggested the possibility of performing similar analysis on tuberculosis patient granuloma samples and provided the impetus for the translational studies reported here. Such analyses have the potential to provide insight into cellular interactions and functions of infected cells within nodal granulomas [12], as well as identify circulating Mtb strains as protocols improve and coverage of the bacterial genome increases.

BLAST analysis of the MTB reads detected in the cell co-culture system uniformly identified Mtb as the top match in every case. This was obtained in only 3 of the 9 clinical samples discussed here. Nevertheless, we believe that the bacterial transcripts detected in the 6 clinical samples that did not match Mtb using BLASTn, are also Mtb in origin for several reasons. First, Mtb infection was confirmed in all these patient samples using acid fast microscopy or GeneXpert analysis. Second, the sequences were associated with host cell transcripts indicating intracellular infection, characteristic of Mtb infection. Third, the sequence variations identified in this study, by definition, decrease the sequence homology to the reference genome, and the sequence variations themselves appeared to be a continuum ranging from low to high frequency amongst the patient samples. And, fourth, the characteristic repeated sequence variations identified in the infecting bacterial rrs and rrl transcripts, are repeated among many of the patient samples, differing primarily in the frequency with which they are detected. Those samples in which the sequence variations in the detected FASTQ reads were fewest were the ones that matched the Mtb using BLASTn, yet they still exhibited some of the same sequence variations repeated in the other 6 samples. Nevertheless, it is important to acknowledge the possibility that the granuloma FNAs may harbor other infecting bacteria, in addition to the Mtb, and that the bacterial reads we are analyzing may not arise solely from Mtb.

In the cell co-culture experiments, there was a high degree of nucleotide sequence deviation in the detected rrs or rrl RNAs when compared to the reference Mtb genome. Variation is to be expected, since the default cutoff we used in the Cell Ranger transcriptome assembly was 10 mismatches per read (approximately 150 nucleotides). This accommodates the error prone processes of library generation and sequencing.

Indeed, the expected random single nucleotide substitutions were detectable in most individual transcript reads. Approximately 58% of the reads from the co-culture experiments contained at least 1 nucleotide substitution, approximately 17% of the reads contained multiple sequence variations. Even higher rates of transcript sequence variation were observed in the clinical samples. Over 90% of the detected reads in most of the clinical samples contained at least one sequence variation and many reads contained multiple variations.

The number of detected Mtb reads differed significantly amongst the patient samples. Detection of bacterial reads was scant in samples FNA 17, FNA 18 and FNA 22. Whereas some samples, such as FNA 0, FNA 4, FNA 10, FNA 13, FNA 20 and FNA 21 exhibited high levels of detected bacterial transcripts. We cannot say, but it is tempting to speculate that this might reflect the relative bacterial load in these patients. If that is so, then we might be able to stratify our patient samples into high bacillary versus low bacillary load categories, as is informative for when analyzing which immune parameters contributing to Mtb control in granulomas [32]. The degree of the nucleotide sequence variation also differed significantly amongst the patient samples. Some samples, like FNA 20 exhibited fewer variations per read, and other such as FNA 0, FNA 4, FNA 10, FNA 13 and FNA 21 exhibited higher degrees of variation.

We do not know how or why the 10X Genomics pipeline seems to detect bacterial rRNA so readily. Perhaps it reflects the high degree of transcription of these genes. Or perhaps, the secondary structure of the rrs and rrl RNAs can self-prime replication or are particularly susceptible to priming by the UMI or barcode regions of the poly-T primers. It is even possible that the modified nucleotides that are common in rRNA, miscode occasionally when replicated and amplified in the 10X Genomics pipeline. “Hard to map regions” and contaminant DNA are known causes of sequence variation in DNA sequencing [33], and conceptually similar problems apply here.

The sequence variations in the bacterial rrs and rrl transcripts did not appear to be random, but rather appeared to be reproduced transcription nucleotide sequence variations common to multiple infecting bacteria. Many of the sequence changes appeared in multiple transcripts, originating from multiple individual bacteria, and in bacteria from different patients obtained years apart. While we observed the anticipated error/nucleotide substitutions in 10X pipeline, they did not appear to create the repeated/retained nucleotide variations that were observed.

We attempted to compare reads from highly transcribed genes of the human host cells, to see if a similar pattern of repeated transcript nucleotide variation was discernable. However, none of our samples had sufficient coverage of human rRNA genes to make this comparison. As an alternative, we compared mitochondrial gene transcript reads where we could find sufficient overlap for comparison, recognizing that the mitochondrial transcripts would not exhibit the same degree of secondary structure or base modification as rRNA. Admittedly, this is an inadequate “apples to oranges” comparison. We observed random errors expected from the 10X genomics pipeline. We also observed a second class of sequence error that was repeated and present if every overlapping read of the mitochondrial gene. We hypothesize this second class of sequence variation, which was uniformly repeated but much less frequent compared to the variations detected in the Mtb reads, were due to actual single nucleotide polymorphisms if the patient mitochondrial genes, but we have no evidence to support this supposition. The fact that this second class of sequence deviation in host cell RNA was omnipresent, and detected in every overlapping MT gene read suggests that it may be a different phenomenon that we detect in the bacterial rRNA reads, which are highly repeated but not necessarily detected in every read from a give patient. However, we do not believe we can conclude this one way or the other.

If one accepts the possibility that the repeated sequence variations seen in the bacterial transcripts actually reflect the transcript sequence, then the data might imply that some selective pressure results in preferred sequence alterations in certain regions of the rrs and rrl RNAs. If such pressure selects for changes that result in functional ribosomal RNA, then perhaps this process of the generation of RNA sequence variation could function to provide short term, evolution mimicking, advantage to Mtb growing under stress.

In conclusion, we have attempted to describe the unexpectedly high degree of sequence variation in the bacterial RNA transcripts we detected. We show that scRNA-seq analysis of nodal human tuberculosis FNA samples is achievable in a resource-limited setting, not requiring refrigerated centrifuges, culture hoods, etc. We also have shown that, unintendedly, scRNA-seq analysis captures RNA from infecting Mtb and can identify the individual cells that harbor the intracellular pathogen [12]. It is disappointing that the coverage of the detected transcripts is insufficient to identify the lineage of the infecting Mtb. However, the efficiency and depth of coverage obtainable with single cell RNA analysis is improving and may soon achieve this goal via analysis of more lineage-differentiating transcripts than rrs and rrl. It is heartening to show that for some FNA samples there is sufficient coverage of rrs and rrl genes to identify mutations associated with drug resistance. When we examined FNA 20 for rrs and rrl transcript variations that coincided with known drug resistance markers [34] we found very few that aligned, even though this patient’s disease was noted as recurrent. The specifics of this patient’s previous medical history are confidential [12].

We document, along with the random sequence variations characteristic of stochastic sequencing errors, non-random repeated nucleotide changes the Mtb rRNA. It is possible that these arise from a stochastic process originating within the 10X Genomics pipeline, due to hard to sequence characteristics of the rRNA such as secondary structure or rRNA nucleotide modification. If so, then the detected variations would not reflect the actual bacterial rRNA transcript sequences. However, if the non-random, repeated nucleotide variations detected in the rRNA reads reflect actual rRNA sequence variation within the bacterium, and if they result in functional ribosomal rRNAs, then one might speculate they could amount to an additional level of epigenetic microbial variation in competitive or challenging environments.

## Materials and methods

### Reagents

GFP H37Ra was a gift from Prasit Palittapongarnpim, Department of Microbiology, Mahidol University, Thailand [22, 23]. Middlebrook 7H9 broth and OADC medium supplement were obtained from BD Biosciences (cat. # 271310 and 211886, respectively; San Jose, CA). Middlebrook 7H10 agar and ADC were obtained from Remel (cat. # R453982 and 705565, respectively; Lenexa, KS). Glycerol was obtained from Acros Organics (cat. # 41098-5000; Fair Lawn, NJ) and Tween 80 from MP Biomedicals, Inc. (cat. # 103170; Santa Ana, CA). THP-1 cells were obtained from ATCC (Cat#TIB-202). HyClone™ RPMI 1640, kanamycin sulfate, Corning™ Accutase™ detachment solution and phorbol 12-myristate 13-acetate (PMA) were obtained from Fisher Scientific (cat. # SH30011.03, BP906-5, MT25058CI, and BP685-1, respectively). Fetal bovine serum was purchased from Atlanta Biologicals (cat. # S11150). BD Horizon™ Fixable Viability Stain 450 was obtained from BD Biosciences (cat. # 562241). Fluorescent Dye 405-I Phalloidin was purchase from Abnova™ (cat. # U0278).

### Bacterial culture

For Mtb *gfp*, bacteria were initially grown on 7H10 supplemented with OADC and kanamycin (KAN; 50 μg/mL). A starter culture of around 10 mL from an isolated colony was grown in 7H9 supplemented 10% ADC, 0.2% glycerol (v/v), 0.05% tween 80 (v/v) and KAN (50 μg/mL) at 37°C until it reached mid-to-late log phase, measured by OD_600_. An aliquot of the starter culture was added to ∼100 mL of 7H9 with KAN (50 μg/mL) and grown to log phase, as monitored by OD_600_.

### Cell culture

THP-1 cells were grown in RPMI 1640 supplemented with 10% FBS and KAN (50 μg/mL) at 37°C, 5% CO_2_.

### GFP Mtb-THP-1 cell co-culture

THP-1 cells were pre-incubated overnight in PMA (20 ng/mL) at 500,000 cells/well in order to generate differentiated macrophages. Medium was replaced for THP-1 cells one hour prior to the addition of DHIV-mCherry and Mtb *gfp*. Preliminary bacterial concentration experiments were performed in THP-1 cells prior to co-culture experiments. An MOI of 2:1 was chosen due to its ability to achieve a high degree of Mtb infection (∼20%) with minimal toxicity to the cells following 24-hour co-incubation. Mtb was added to THP-1 cells and incubated for or Mtb 5 days followed by preparation for analysis by flow cytometry and cell imaging or scRNA-seq analysis.

### Flow cytometry

Adherent THP-1 cells were incubated with Accutase™ for 15 minutes at 37°C. THP-1 cells were transferred to 5-mL tubes and washed with PBS. Cells were resuspended in BD Horizon™ Fixable Viability Stain 450 (0.25 μg/mL) and incubated at 4°C for 30 minutes. Cells were then fixed in 2% formaldehyde for 30 minutes at 4°C. Cells were analyzed using a FACS Canto. Percent infection by DHIV was quantified as a subset of the live population (FSC/V450/50-). Gates for infection were set according to the uninfected “mock” THP-1 cell controls. Three independent biological replicates were completed for all treatment conditions, each in triplicate wells per experiment.

Population analysis was then done using FlowJoTM v10.7 [35], to assess if infection levels and cell viability were consistent similar in all replicates. The Flow Cytometry figures are representative plots obtained from one of the replicates. Minimum of 30,000 events were collected per experiment.

### Cell imaging

Following Accutase™ removal of adherent THP-1 cells, THP-1 cells were washed with PBS fixed with 2% formaldehyde at 4°C for 30 minutes. Cells were resuspended in 100 μL of a 1:1000 dilution of Fluorescent Dye 405-I Phalloidin in PBS and incubated for 15 minutes at room temperature. The images were acquired on a Nikon A1 confocal microscope using a 60x oil lens. Images were processed using Fiji [36].

### Preservation of fine needle aspirates for genomic and transcriptomic analysis

Under the University of Papua New Guinea School of Medicine-Medical Research Council approved protocol, patients presenting at the CPHL TB Clinic, and tentatively diagnosed with LNTB, upon giving informed consent, were subjected to the standard diagnostic protocol, which includes FNA of enlarged (>1.0 cm) lymph nodes. This aspirate goes for microscopy and for GeneXpert analysis as part of the patient assessment process. Aspirate from one pass of the granuloma dedicated to this study was washed directly into ice-cold RPMI buffer containing 0.2% fetal bovine serum, in a heparinized tube. The samples were taken to the adjacent pathology lab where they are pelleted for 5 min. and suspended on ice in NH_4_Cl lysis solution [37] to lyse contaminating erythrocytes. After a maximum of 5 min with occasional gentle mixing on ice, and observation of the depletion of obvious erythrocytes, 1 mL of Accutase™ was added directly to the lysis buffer for a maximum of an additional 3 min., again with occasional gentle mixing and observation of the dissolution of obvious tissue clots in the solution. The sample volume was expanded with ice cold RPMI buffer and the cells were pelleted again. The pelleted cells were gently suspended in 200 µl preservation buffer to which 1 mL of ice-cold methanol is slowly added with mixing. The de-identified samples are kept on ice packs for transportation and analysis by PCR, WGS and scRNA-seq.

### Single-cell RNA-sequencing

scRNA-seq was performed on single-cell suspensions using either the 10X Genomics Chromium to prepare cDNA sequencing libraries as described by Brady et al. [27].

Samples were processed using the Chromium Single Cell 3′ V3 Kit (10X Genomics, Cat. # 1000075) using whole cells fixed in 80% methanol. Single cells were diluted to a target of 1000 cell/μL in 1× PBS (whole cells) or 1× PBS + 1.0% BSA + 0.2 U/μL RiboLock RNase Inhibitor to generate GEM’s prepared at a target of 10000 cells per sample. Barcoding, reverse transcription, and library preparation were performed according to manufacturer instructions. 10X Genomics generated cDNA libraries were sequenced on Illumina HiSeq 2500 or NovaSeq 6000 instruments using 150 cycle paired-end sequencing at a depth of 10K reads per cell. scRNA-seq was performed at the High Throughput Genomics Core at Huntsman Cancer Institute (HCI) of the University of Utah.

For analytical procedures, the 10X Genomics Cell Ranger Single Cell software pipeline [13] was deployed to produce alignments and counts, utilizing the prescribed default parameters. The human genomic reference was GRCh38 and the Mtb genome H37Rv was used for alignment. For quality management and further analytical exploration, Seurat (4.1.0) was utilized. Doublets were identified with DoubletFinder, cells were excluded based on having less than 100 genes and an excess of 25% mitochondrial genes. Mitochondrial genes, were filtered out but every cell that contained Mtb genes was retained. Dimensionality was reduced and scaled via SCTransformation (0.3.5) using the Gamma-Poisson generalized linear model (glmGamPoi, 1.4.0) methodology at default resolution. Automated categorization of cells was performed using SingleR (1.6.1) [38–51]. Statistics within Seurat pipelines were generated with FindAllMarkers or FindMarkers [51].

### Single-cell RNA-sequencing

scRNA-seq was performed on single-cell suspensions using 10X Genomics Chromium to prepare cDNA sequencing libraries as described by Brady et al. [13, 44]. The samples were processed using the Chromium Single Cell 3′ V3 Kit (10X Genomics, Cat. # 1000075) using whole cells fixed in 80% methanol. Single cells were diluted to a target of 1000 cell/μL in 1× PBS (whole cells) or 1× PBS + 1.0% BSA + 0.2 U/μL RiboLock RNase Inhibitor to generate GEM’s prepared at a target of 10000 cells per sample. Barcoding, reverse transcription, and library preparation were performed according to manufacturer instructions. 10X Genomics generated cDNA libraries will be sequenced on NovaSeq 6000 instruments using 150 cycle paired-end sequencing at a depth of 10K reads per cell. The scRNA-seq was performed at the High Throughput Genomics Core at Huntsman Cancer Institute of the University of Utah.

For analytical procedures, the 10X Genomics Cell Ranger Single Cell software pipeline [13] is deployed to produce alignments and counts, utilizing the prescribed default parameters. The genomic references used for alignment were the human (hg38), the H37Rv Mtb (NC_00096.3:1) and HIV-1 (NC_001802.1). For quality management and further analytical exploration, Seurat (4.1.0) was utilized. Doublets were identified with DoubletFinder, cells were excluded based on having less than 100 genes/features and an excess of 25% mitochondrial genes. Mitochondrial genes were filtered out but every cell that contained Mtb genes was retained. Dimensionality was reduced and scaled via SCTransformation (0.3.5) using the Gamma-Poisson generalized linear model (glmGamPoi, 1.4.0) methodology at default resolution or less. Automated categorization of cells was performed using SingleR (1.6.1). Statistics within Seurat pipelines were generated with FindAllMarkers or FindMarkers which utilizes a Wilcoxon rank sum test [51].

## Statistical analysis

Unpaired t tests were used to determine statistical significance across infection culture conditions of THP-1 from three independent experiments. Significance was determined and the level recorded if the p-value was less than 0.05.

The pairwise TTests function from Scran was used to determine statistically significant differential expression of genes between groups. This was performed for all comparison sets. Other default statistical standards were adopted from the various software recommendations during data analyses unless otherwise specified [65].

## Acknowledgments

This work was supported by a University of Utah seed grant (LRB) and an ALSAM Foundation Grant (LRB & PJM). AFC recognizes support from K08 AI139339 and the University of Utah Department of Pathology start-up funds. Mr. A.L. Lim was partially supported during this work by the grant U01TW008163. The authors also wish to acknowledge Professor Nakapi Tefuarani, Dean UPNG School of Medicine for his enduring support, and Dr. Rodney Itaki and Dr. Evelyn Lavu (decd), UPNG School of Medicine, for early assistance in obtaining our IRB. The authors sincerely acknowledge the multiple contributions of Dr. Erica C. Larson, Dept. Microbiology and Mol. Genetics, Univ. Pittsburgh, who headed the develop the THP-1-Mtb co-culture system while training at Univ. Utah., and who made many constructive contributions to the development of this manuscript. This work was supported by the University of Utah HCI Flow Cytometry Facility and Metabolomics core facilities, and the HSC High Throughput Genomics core facility. The help of Chris Stubben in bio-statistical analysis is gratefully acknowledged: "Research reported in this publication utilized the High-Throughput Genomics and Cancer Bioinformatics Shared Resource at Huntsman Cancer Institute at the University of Utah and was supported by the National Cancer Institute of the National Institutes of Health under Award Number P30CA042014. The content is solely the responsibility of the authors and does not necessarily represent the official views of the NIH."

