## Supplementary Information for "Description of Bacterial RNA Transcripts Detected in *Mycobacterium tuberculosis* – Infected Cells from Peripheral Human Granulomas using Single Cell RNA Sequencing"

**Supplemental Figures**

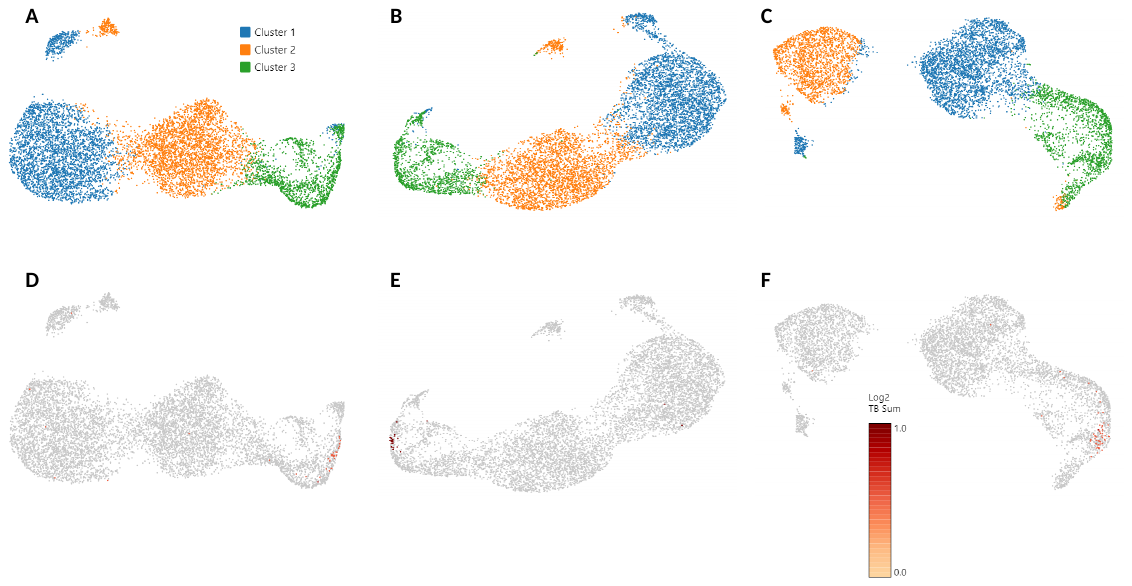

**Fig S-1 A-C)** **K means clustering, K=3, of THP-1/Mtb co-cultures analyzed 5 days after inoculation of 0.5X10^6^ cells with H37Ra.** THP-1 cells were differentiated with PMA for 24 h prior to infection and then co-incubated with GFP H37Ra (MOI 2:1). At K=3 clustering Cluster #3 closely matches the percentage of GFP expressing cells detected in replicate cultures by Flow Cytometry. **D-F)** Feature plot showing any cell containing a Mtb transcript. Mtb containing cells were almost exclusively found in Cluster #3. **A)** Experiment 14800X3, percentage of cells in cluster 3 was approximately 15%, percentage GFP positive by flow cytometry approximately 15%. **B)** Experiment 14800X4 technical repeat, percentage of cells in Cluster #3 was approximately 17%. **C)** Exp. 14800X5, technical repeat of 14800X6 presented in Fig.2, percentage of cells in Cluster #3 was approximately 24%, percentage GFP positive by flow cytometry approximately 24%.
